# Persistent Decision-Making in Mice, Monkeys, and Humans

**DOI:** 10.1101/2024.05.07.592970

**Authors:** Veldon-James Laurie, Akram Shourkeshti, Cathy S Chen, Alexander B Herman, Nicola M Grissom, R. Becket Ebitz

**Affiliations:** University of Montreal; University of Minnesota

## Abstract

Humans have the capacity to persist in behavioural policies, even in challenging environments that lack immediate reward. Persistence is the scaffold on which many higher executive functions are built. However, it remains unclear whether humans are uniquely persistent or, instead, if this capacity is widely conserved across species. To address this question, we compared humans with mice and monkeys in harmonised versions of a dynamic decision-making task. The task encouraged all species to strike a balance between persistently exploiting one policy and exploring alternative policies that could become better at any moment. Although all three species had similar strategies, we found that both primate species—humans and monkeys—were able to persist in exploitation for longer than the mice. The similarities in persistence patterns in humans and monkeys, as opposed to mice, may be related to the various ecological, neurobiological, or cognitive factors that differ systematically between these species.

## INTRODUCTION

Decision-making in uncertain environments requires the flexibility to balance exploiting previously rewarded options with exploring alternatives that could be better. This flexibility is an important part of human intelligence (Rich and Gureckis, 2018; Knox et al., 2012; Wilson et al., 2021; Yan et al., 2023), yet it is also fragile and often compromised by various neurobiological conditions (Kaske et al., 2023; Verdejo-García et al., 2006; Tolin et al., 2009; Blanco et al., 2013; Teng et al., 2016; Mäntylä et al., 2012). Because evolution tends to canalise phenotypes over time (Waddington, 1942; Siegal and Bergman, 2002), this fragility could suggest that the human solution to the explore-exploit dilemma may have evolved recently. However, in part because of the difficulty of harmonising tasks and data collection across species, we do not know how human exploratory decision-making compares with other species.

Comparative studies of exploratory decision-making are particularly urgent as non-human animal models are often used to model human cognitive functions. Due to the development of optogenetics and other circuit-level techniques in mice (Boyden et al., 2005), there has been an explosion in cognitive function research in this species (Ellenbroek and Youn, 2016) and many influential studies of decision-making under uncertainty have focused on rodent models (Saddoris et al., 2015; Groman et al., 2016; Bari et al., 2019; Izquierdo et al., 2019; Soltani and Izquierdo, 2019; Chen et al., 2021a, 2021b; Grossman et al., 2022; Iyer et al., 2022). However, it remains unclear whether mice and other rodents solve these tasks in a way that could justifies their use as a model species for humans (Manger et al., 2008; Stevenson et al., 2018; Woo et al., 2023). Further, the limited amount of comparative work that exists already suggests that there will be differences between species in these tasks (Trepka et al., 2021; Woo et al., 2023), though no study has yet directly compared mice to humans.

Here, we compared human exploratory decision-making with two commonly used animal models in cognitive neuroscience: Mus musculus (the mouse) and Macaca mulatta (the rhesus monkey). These species deviated from the human lineage at radically different times (monkey: ∼24 million years ago (Disotell and Tosi, 2007; Gibbs et al., 2007); mice: ∼90 million years ago (Ernst and Carvunis, 2018)). To identify the similarities and differences between species, we had each species perform “harmonised” versions of a classic explore/exploit task: a restless multi-armed bandit where they made a series of choices between targets with unpredictable rewards. Tasks were “harmonised” across species in the sense that rewards were tied to spatial location and key parameters of the reward schedules were matched, though each was administered via species-typical experimental equipment. Although all three species showed similar behavioural signatures of exploration and exploitation, there were differences in how the species switched between targets. Computational modelling revealed that the key difference between mice (who switched frequently) and monkeys and humans (who did not) lay in the primates’ capacity to persistently exploit targets for longer than mice. Three control experiments in humans ruled out several low-level explanations for these species’ differences. Together, these results suggest that the primate lineage may have only recently evolved its remarkable capacity to persist in exploitative states.

## RESULTS

In **Experiment 1**, mice (N=32; average of 7.97 sessions each; 255 sessions; 70,478 total trials), monkeys (N=5; average of 18.6 sessions; 93 sessions; 57,878 total trials) and humans (N=258; 1 session each; 258 sessions; 77,400 total trials) performed comparable spatial restless k-armed bandit tasks (**Figure 1A**). Choice options (targets) offered a probability of reward which changed slowly and independently over time (**Figure 1B**). The task encouraged participants to both exploit rewarding targets and explore new targets to learn about potential rewards. Mice indicated their choices via nose pokes, monkeys via saccadic eye movements, and humans with a computer mouse (**Figure 1A**). Effects of differences in the timing of the task, the number of targets and the response modality were estimated via 3 control experiments in humans (**see Methods**; **Experiment 2, 3, 4**).

**Figure 1.**
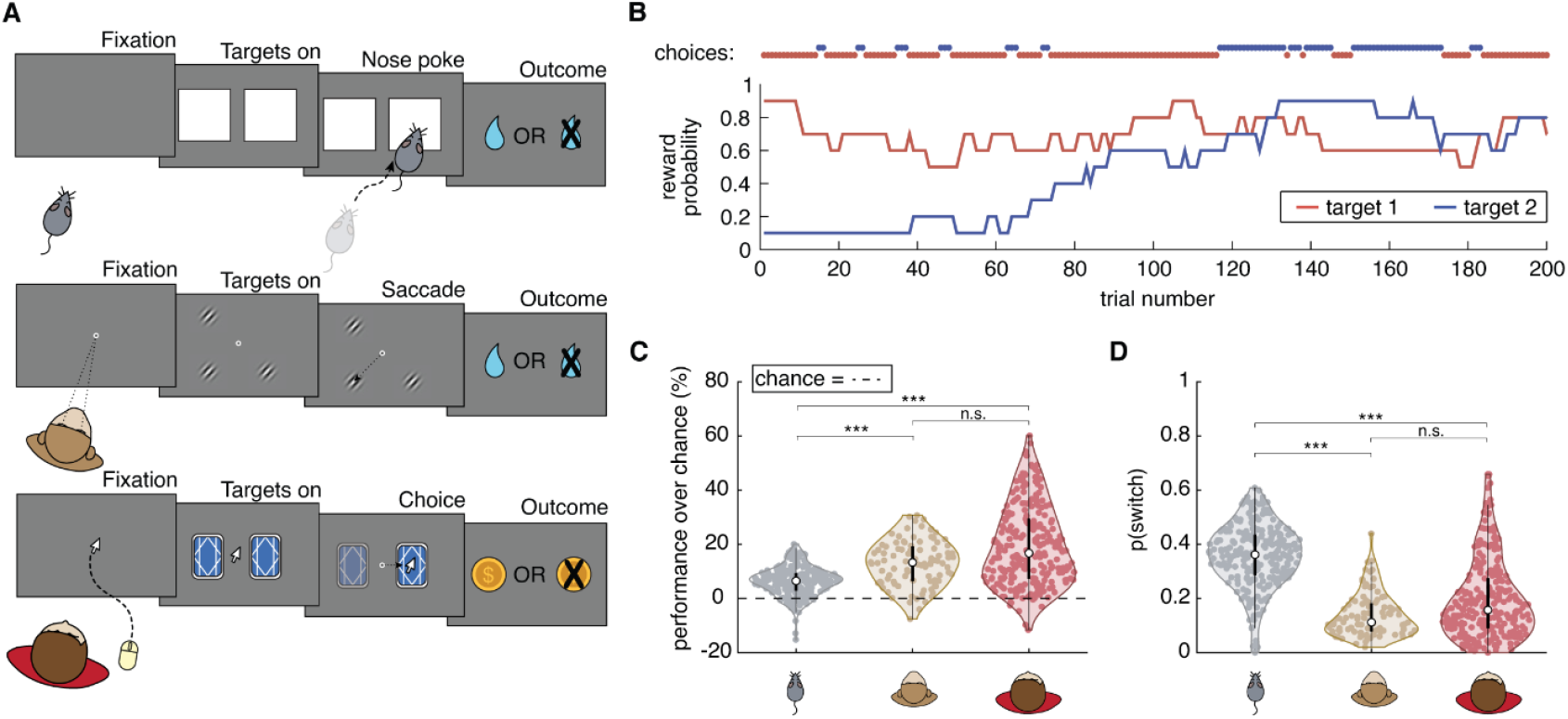
Task design and behaviour across species. **A**) Schematic representation of the bandit task in each species (mice = top, monkeys = middle, humans = bottom). **B**) Example reward schedule, including 200 trials from one session with one human. The reward probabilities of the 2 targets (blue and red) walk randomly, independently across trials. The humans’ choices are illustrated as coloured dots along the top. **C**) Percentage of reward relative to chance in all species. Thick black lines = IQR, thin = whiskers, open circle = median. Black dotted line = chance performance. **D**) Probability of switching targets between species. Same conventions as C. * indicates α = 0.05; ** indicates α = 0.001; and *** indicates α = 0.0001, corrected for multiple comparisons.

There were differences in performance between species: humans and monkeys were more likely to get rewards compared to mice (normalised difference from chance; **Figure 1C**; humans-mice: p > 0.0001, b = -13.04, 95% CI = [-17.78, -8.29], t(511) = -5.40; monkeys-mice: p < 0.0001, b = – 7.65, 95% CI = [–10.94, –4.37], t(344) = -4.59). Humans and monkeys performed equivalently (humans-monkeys: p = 0.24, b = -6.66, 95% CI = [-17.77, 4.46], t(349) = -1.18). Humans and monkeys switched less often than mice (**Figure 1D**; humans-mice: p > 0.0001, b = 0.19, 95% CI = [0.12, 0.20], t(511) = 7.21; monkeys-mice: p < 0.0001, b = 0.26, 95% CI = [0.18, 0.34], t(344) = 6.17). There were no differences in switch probability between humans and monkeys (**Figure 1D**; p = 0.33, b = -0.056, 95% CI = [-0.17, 0.057], t(349) = -0.98). There were no differences in performance or the probability of switching (or additional measures, like the probability of exploration) with either self-reported gender in humans or sex in mice (**Figure S1**; all monkeys were male). In sum, species differences in both performance and switching were driven by differences between the primate species and the mice.

### Switching dynamics and exploratory behaviour

Switching happens for multiple reasons in this task (Ebitz et al., 2018; Chen et al., 2021b). Sometimes animals switch options because they are engaging in rapid trial and error sampling. Other times they switch because the option they have been choosing is no longer rewarding. To differentiate these forms of switching, we fit a mixture model to the distribution of inter-switch intervals in each species (**Figure 2A**; **see Supplemental Methods**).

**Figure 2.**
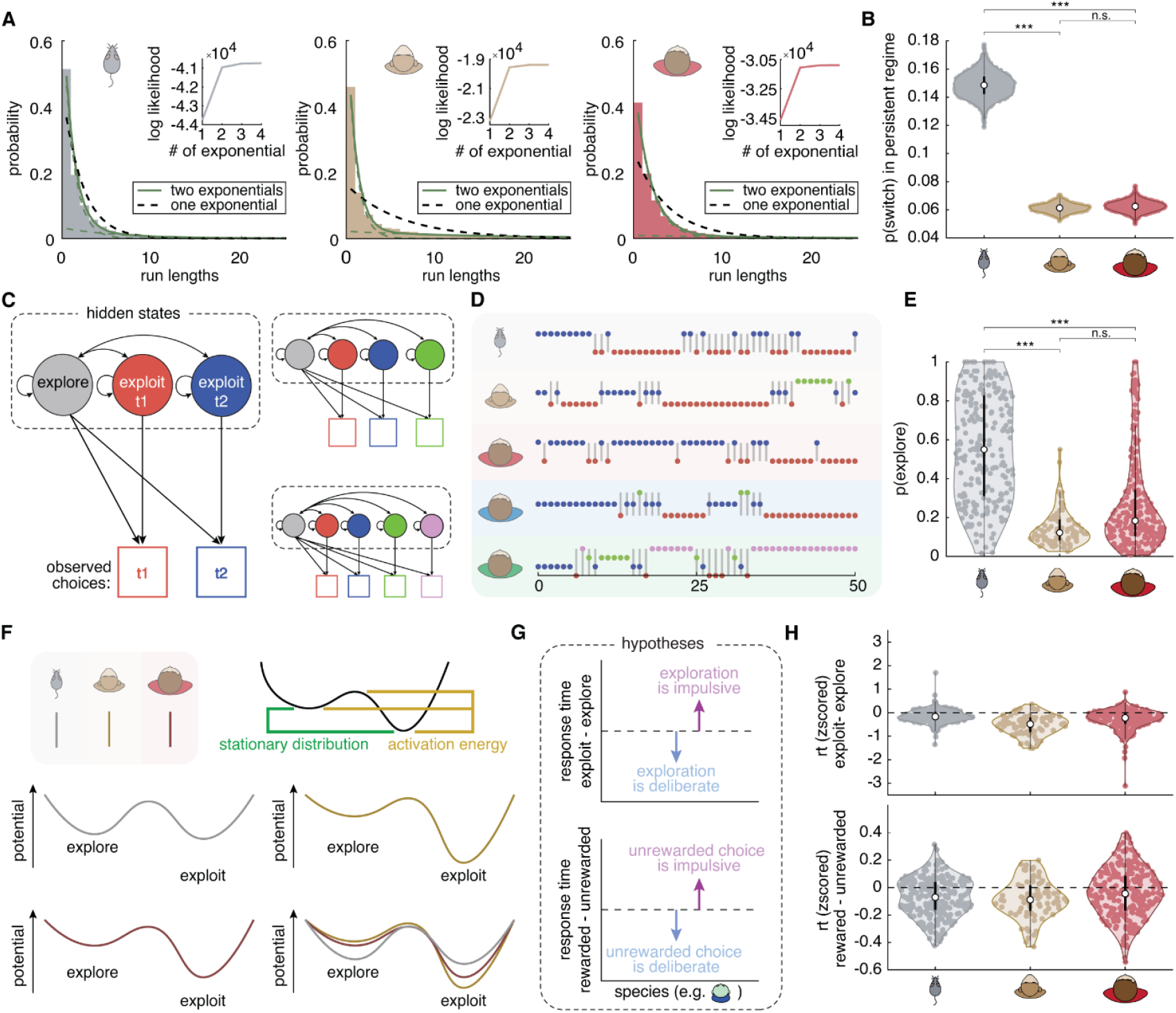
Different patterns of switching and exploration across species. **A**) Distributions of the number of trials between switch decisions (“run lengths”) in mice, monkeys and humans. If there was a fixed probability of switching, run lengths would be exponentially distributed (black dotted line). A mixture of two exponential distributions (green line) suggests 2 distinct probabilities of switching. (Inset) Log likelihoods for different mixture models containing 1 to 4 exponential distributions in each species. **B**) Bootstrapped estimates of switch probability for the slow-switching distribution (the “persistent regime”) across species. Thick lines = IQR, thin = whiskers, open circle = median. **C**) Hidden Markov models (HMMs) were used to infer goal states for each choice. The model included one persistent state for each target (‘‘exploit’’) and one state where any of the targets could be chosen (“explore”). Right) HMMs were extended for different numbers of targets. **D**) Example choice sequences for mice, monkeys and humans. Circles represent the chosen target; grey lines highlight explore choices. **E**) Probability of exploration across species, as inferred by the HMM. Same conventions as B. **F**) Fitting the HMM identifies the equations that describe the dynamics of exploration and exploitation, meaning the rate at which participants explore, exploit, and switch between states. Left) Measures of these equations, namely their stationary distributions (Boltzmann, 1868) and activation energies (Arrhenius, 1889) can be used to derive the energetic landscape of the states. Middle) Average state dynamic landscapes. Right) State dynamic landscapes for all species overlaid **G**) Differences in response time (RT) reveal which choices are fast and impulsive or slow and deliberative. Top: a cartoon illustrating RT differences under the hypothesis that exploration is more deliberative than exploitation (blue) or that exploration is more impulsive (purple). Bottom: same for rewarded and unrewarded trials. **H**) Z-scored RT differences across all three species. Top: difference across HMM-defined exploration and exploitation. Bottom: between rewarded and unrewarded trials. Thick black lines = IQR, white circle = median. * Indicates α = 0.05; ** indicates α = 0.001; and *** indicates α = 0.0001, corrected for multiple comparisons.

The behaviour of all species was best described as a mixture of two switching modes (**Figure 2A, Supplementary Table 1**). All participants sometimes switched between targets at a fast pace (“switching regime”) and sometimes stuck to choosing one target repeatedly (“persistent regime”) and the two regimes occurred equally often across species (**Supplementary Table 2**). Humans and monkeys differed from mice in their average switching probability during the persistent regime (**Figure 2B**; humans-mice: p < 0.0001, b = 0.081, 95% CI = [0.056, 0.11], t(500) = 6.32; monkeys-mice: p < 0.0001, b = 0.16, 95% CI = [0.11, 0.22], t(341) = 5.77). Again, there were no differences between primates (humans-monkeys: p = 0.29, b = -0.056, 95% CI = [-0.16, 0.047], t(341) = - 1.063). This suggest that species differences in switching (**Figure 1B**) were largely driven by the primates’ increased tendency to persist in the persistent regime, compared to mice. However, humans also differed from both monkeys and mice in the average switch probability during the switching regime (humans-mice: p < 0.0001, b = 0.19, 95% CI = [0.14, 0.24], t(500) = 6.94; humans-monkeys: p < 0.017, b = 0.15, 95% CI = [0.034, 0.27], t(341) = 2.53). Monkeys did not differ from mice here (p = 0.28, b = 0.054, 95% CI = [-0.044, 0.15], t(341) = 1.083). This could suggest that humans use a different strategy for exploratory sampling than either monkeys or mice, even as monkeys share the human increase in persistence compared to mice.

In order to determine why primates were more persistent than mice, we categorised individual choices based on the underlying reason for those decisions with Hidden Markov Models (HMMs; **Figure 2C-D**; **see Supplemental Methods**). This method probabilistically infers whether choices are due to a state of exploration (trial- and-error sampling) or a state of exploitation of a single option (Ebitz et al., 2018, 2019; Chen et al., 2021b; Kaske et al., 2023). Based on the HMM labels, the probability of exploring differed between both humans and monkeys, compared to mice (**Figure 2E**; humans-mice: p < 0.0001, b = 0.29, 95% CI = [0.22, 0.35], t(479) = 8.82; monkeys-mice: p < 0.0001, b = 0.50, 95% CI = [0.33, 0.67], t(327) = 5.82). There were no differences within primates (humans-monkeys: p = 0.22, b = -0.13, 95% CI = [-0.33, 0.076], t(334) = -1.24). These results suggest that primates were less exploratory than mice.

We next asked if there were species differences in the temporal dynamics or stability of exploration and exploitation. First, we simply compared the fitted parameters of the HMMs (Ebitz et al., 2018, 2019; Chen et al., 2021b), and found species differences in the likelihood of staying in exploitation (exploit-to-exploit transition probability: humans-mice: p < 0.0001, b = -0.092, 95% CI = [-0.12, -0.067], t(479) = -6.98; humans-monkeys: p = 0.32, b = 0.055, 95% CI = [-0.054, 0.16], t(334) = 1.00; monkeys-mice: p < 0.0001, b = -0.16, 95% CI = [-0.22, -0.095], t(327) = -5.00). Specifically, mice were less likely to stay in exploitation than either humans or monkeys (mean mice: 0.77 ± 0.19 STD across sessions, monkeys: 0.92 ± 0.046 STD, humans: 0.86 ± 0.13 STD).

Next, we performed certain mathematical analyses of the HMM solutions (**Figure 2F**; see **Supplemental Methods**). Briefly, the fitted parameters of the HMM represent a system of equations that describe the dynamics of exploration and exploitation in each session. Through analysing these equations, we can infer (1) the difference in potential energy between states (a measure of the difference in the stability), and (2) the activation energy needed to transition from exploitation to exploration (a measure of the stability of exploitation). In mice, the potential energy of exploration and exploitation were equivalent (mean difference in energy = 0.19 ± 1.55 STD across sessions), suggesting that each state was equivalently easy to occupy. By contrast, exploitation was a deeper, more energetically stable state than exploration in both primate species (monkeys: -1.87 ± 0.72 STD; humans: -1.30 ± 1.61 STD). Humans and monkeys were both significantly different than mice (humans-mice: p < 0.0001, b = 1.47, 95% CI = [1.053, 1.88], t(479) = 6.95; monkeys-mice: p < 0.0001, b = 2.29, 95% CI = [1.30, 3.27], t(327) = 4.56), but not different from each other (humans-monkeys: p = 0.38, b = -0.58, 95% CI = [-1.87, 0.72], t(334) = -0.88). The amount of energy required to end exploitation was lower in mice than humans, whereas humans and monkeys did not differ (differences in activation energy in humans-mice: p < 0.0001, b = -0.74, 95% CI = [-1.08, -0.40], t(479) = -4.30; humans-monkeys: p = 0.84, b = 0.13, 95% CI = [-1.13, 1.39], t(334) = 0.20; mean across species in mice = 1.77 ± 1.39 STD, monkeys = 2.69 ± 0.57 STD, humans = 2.54 ± 1.53 STD). Slightly more energy was required to end exploitation in monkeys than mice, but this did not survive correction for multiple comparisons (monkeys-mice: p = 0.029, b = -0.083, 95% CI = [-1.58, -0.084], t(327) = -2.19). In short, exploitation was more stable in both primates compared with mice.

### Impulsivity does not explain mouse switching dynamics

It is possible that mice were less persistently exploitative because they made more “impulsive” decisions, increasing the frequency of “exploratory” choices that were not truly exploratory. However, “impulsive” choices would be faster than other kinds of choices—in contrast to genuine exploratory choices, which tend to be slower (Ebitz et al., 2018; Chen et al., 2021b). Therefore, to determine if there were species differences in impulsivity, we analysed the reaction times in the explore/exploit trials (**Figure 2G, top**). To permit comparison across species with differing response modalities, we focused on how response time differed across states through calculating a modulated index (see **Methods**). All 3 species were faster for exploitation compared to exploration (modulated index < 0; **Figure 2H - top;** different from zero in mice: p < 0.0001, b = - 0.15, 95% CI = [-0.19, -0.099], t(222) = -6.08, monkeys: p < 0.0002, b = -0.51, 95% CI = [-0.75, -0.26], t(78) = -4.093, humans: p < 0.0001, b = -0.28, 95% CI = [-0.34, -0.23], t(238) = -10.52). This implies that exploration was deliberative, rather than impulsive, in all three species.

Although exploratory choices were not more impulsive in mice as a rule, it remained possible that mice performed worse because of impulsivity. Therefore, we next performed the same response time analysis for rewarded versus unrewarded trials (**Figure 2G, bottom**). We found that all species had slower reaction times on unrewarded trials compared to rewarded trials (**Figure 2H - bottom;** different from zero in mice: p < 0.0001, b = -0.071, 95% CI = [-0.088, -0.054], t(254) = -8.21; monkeys: p < 0.001, b = -0.079, 95% CI = [-0.13, -0.034], t(78) = -3.48; humans: p < 0.0005, b = -0.040, 95% CI = [-0.062, -0.018], t(257) = -3.57), suggesting that incorrect choices were not caused by impulsivity in mice. This indicates that the mice’s strategy, while less rewarding, did not appear to be a more impulsive strategy.

### Manipulating task variables to understand species differences

Species differences in performance, exploration, and persistence, could have been artifacts of the small variations in task design across species. For example, to keep two of the monkeys motivated in the task, the minimum probability of a reward had to be increased from 0.1 to 0.3. As a result, two monkeys experienced reward walks that were slightly richer on average than the other monkeys, mice, and humans. Still, this did not impact the probability of switching (Normal Env. vs Rich Env.: p=0.77, b = -0.013, 95% CI = [-0.099, 0.073], t(89) = -0.29), nor the probability of exploration (monkey Normal Env. vs monkey Rich Env.: p = 0.91, b = 0.0051, 95% CI = [-0.087, 0.097], t(89) = 0.11). These 2 monkeys were essentially identical to the other 3 monkeys whose reward schedule matched the other participants. To determine if species differences were related to other task differences, we looked at the effects of variations in the number of targets (**Experiment 2**), task timing (**Experiment 3**), and differences in response modality (**Experiment 4**) in humans.

Monkeys did a 3-target version of the task, while mice and humans did 2-target versions. Therefore, it is possible that monkeys were more similar to humans because adding a third target (1) improves reward acquisition, (2) reduces switching, and (3) decreases exploration. However, this was not the case. In an online sample of 150 humans (1 session each, 45 000 total trials), we manipulated the number of targets (2, 3, or 4 targets). The likelihood of getting rewards did increase with the number of targets (**Figure 3A;** normalised difference from chance; one-way ANOVA: F_2, 141_ = 16.03, p < 0.0001, S = 144 total sessions), but so did switching, though this was at trend (**Figure 3B;** one-way ANOVA: F_2, 141_ = 2.85, p = 0.061, S = 144 total sessions). Further, increasing the number of targets had no effect on exploration (**Figure 3C;** one-way ANOVA: F_2,132_ = 0.75, p = 0.48; S = 135 total sessions). This implies that changing the number of targets was not a sufficient explanation for differences in either the probability of switching or the probability of exploration between species. Manipulating the number of targets also did not alter the probability of repeating exploitation (i.e. the HMM parameter, one-way ANOVA: F_2, 132_ = 0.53, p = 0.53; S = 135 total sessions; **Figure 3D**), the relative energy of exploration and exploitation (F_2, 132_ = 0.99, p = 0.37), the energy barrier between the states (F_2, 132_ = 2.66, p = 0.074). These results show that simply changing the number of targets cannot fully explain the species difference in persistence.

**Figure 3.**
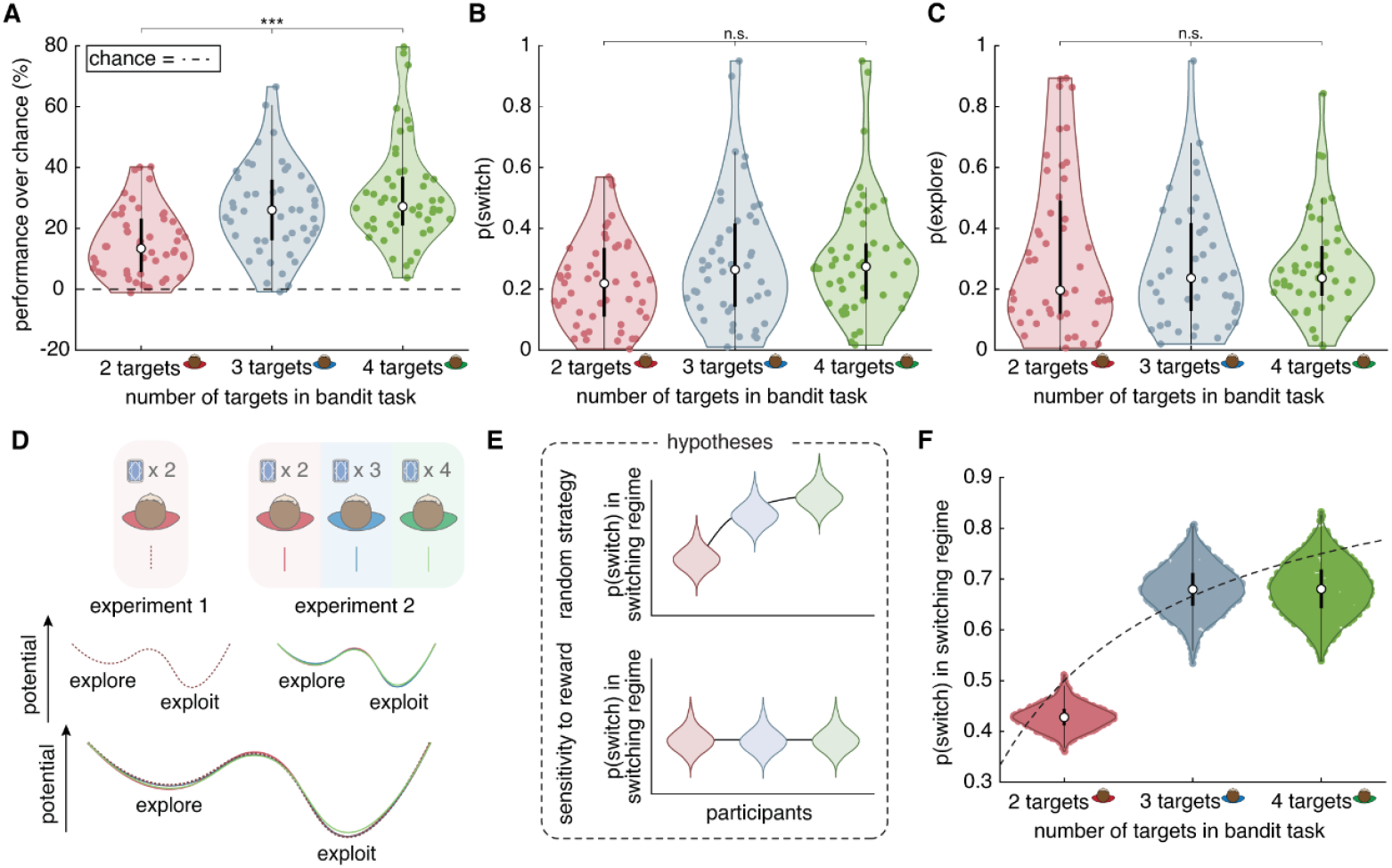
Effects of manipulating the number of targets in humans performing the bandit task (Experiment 2). **A**) Percentage of reward relative to chance by number of targets (2, 3, or 4). Thick black lines = IQR, thin = whiskers, open circle = median **B**) Switch probability by number of targets. **C**) Probability of exploration by number of targets. **D**) State dynamic landscapes for varying numbers of targets (Same conventions as Figure 2F). **E**) Cartoon illustrating predicted relationships between the switching-regime switch probability and the number of targets under the hypothesis of random exploration (top) or reward-dependent exploration (bottom) **F**) Switch probability for the switching regime by number of targets. * Indicates α = 0.05; ** indicates α = 0.001; and *** indicates α = 0.0001.

Although increasing the number of targets did not alter exploration, there was a trend towards more switching. Therefore, we next asked if there were any specific changes in the probability of switching within either the exploitative, persistent regime or in the exploratory switching regime. There was no difference in switching in the persistent regime with the number of targets, again suggesting that species differences in this measure were unlikely to be due to differences in targets (One-way ANOVA: F_2, 139_ = 1.00, p = 0.37; S = 142 total sessions; **see Methods**). There was a moderate increase in switching in the exploratory regime (**Figure 3F;** One-way ANOVA: F_2, 139_ = 10.25, p < 0.0001; S = 142 total sessions). The way that exploratory switching changed with the number of targets implied that humans—like monkeys (Ebitz et al., 2018; Wilson et al., 2021)—used a random form of exploration in this task. In mice, it is not clear whether they also used random exploration because they were not trained on the 3+ target version of the task needed to assess this. Under random exploration, we would expect exploratory switching to increase systematically with the number of targets (**Figure 3E-F, top**; see **Methods**). This is because random choices between a smaller number of targets (i.e. 2) are more likely to repeat (i.e. 50% of the time) than random choices between many targets (4 targets will repeat 25% of the time). By contrast, if exploratory decisions were based on the rewards of the chosen target (i.e., a win-stay, lose-shift strategy), switching would be unaffected by the number of alternatives (**Figure 3E, bottom**). We found that the human pattern of switching followed the prediction of a random exploration strategy (**Figure 3F, Supplementary Table 3**). In sum, while altering the number of targets did not alter persistence in humans, it did suggest that humans used a random sampling strategy for exploration.

Exploratory decision-making is affected by physiological and psychological processes that operate in the time scale of the body, not just in the time scale of trials (Shourkeshti et al., 2023). Further, prior reward information can decay over time, rather than just trials, meaning that it may be harder to recall past rewards when trials take longer. In mice, trials took longer because this species had to physically move from a central point to their desired target and then back to a reward port on every trial. This meant the mouse version of the task was inherently slower than the primates’. To determine if the long trial lengths in mice were a sufficient explanation for their decreased persistence relative to primates, we manipulated trial lengths in humans **Experiment 3**. We did this first by lengthening inter-trial interval times to determine whether the mere passage of time between rewards is sufficient to reduce persistence. In an online sample of 299 human participants (1 session each, 89,699 total trials), we found slight variations in the likelihood of getting rewards across as the inter-trial-interval increased (**Figure 4A;** normalised difference from chance; one-way ANOVA: F_2, 296_ = 3.48, p < 0.05, S = 299 total sessions). However, there was no significant effect of the inter-trial intervals on switching (**Figure 4B;** one-way ANOVA: F_2, 296_ = 0.53, p = 0.59, S = 299 total sessions) or exploration (**Figure 4C**; one-way ANOVA: F_2, 282_ = 0.48, p = 0.62; S = 285 total sessions). Again, we found no systematic differences in performance, switching or exploration by self-reported gender in this experiment (**Figure S2**). These results suggest that the passage of time between decisions did not affect the key behavioural differences between species, making it an unlikely explanation for them. However, because we isolated the timing variable for this control experiment, this result does not rule out the possibility that different response modalities between species could cause differences in persistence.

**Figure 4.**
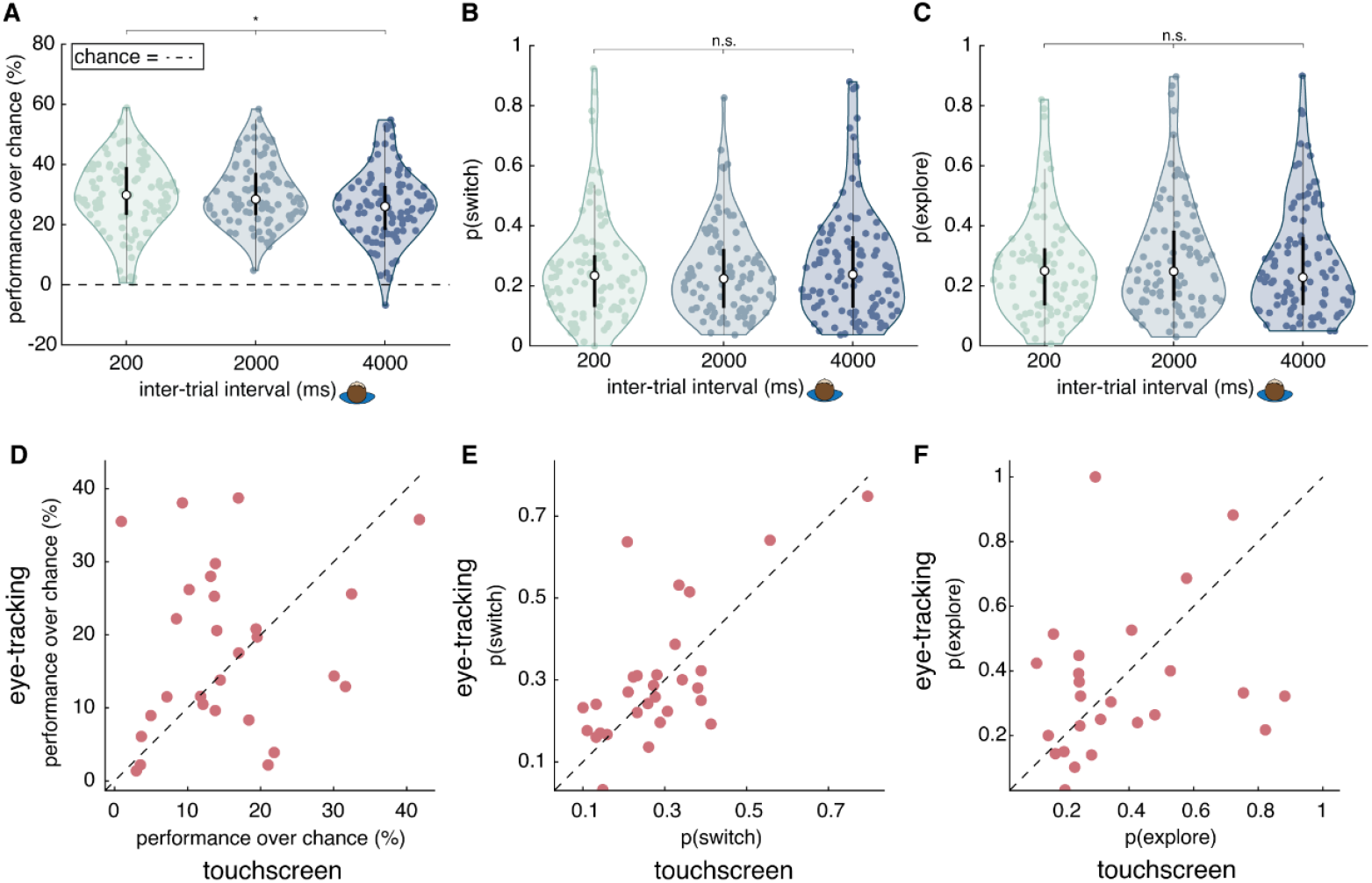
Effects of manipulating the trial length and modality of response in humans performing the bandit task (Experiment 3 and 4). **A)** Percentage of reward relative to chance across varying inter-trial interval times. Thick black lines = IQR, thin = whiskers, open circle = median. **B)** Switch probability by inter-trial interval times. **C)** Probability of exploration by inter-trial interval times. **D)** Within-participant comparison of percentage of reward relative to chance between the touchscreen and the eye tracking response modalities. **E-F)** Same as in D but for probability of (E) switching and, (F) exploration. * Indicates α = 0.05; ** indicates α = 0.001; and *** indicates α = 0.0001.

Given that all species performed bandit tasks using different modalities of response, it remained possible the observed differences were caused by their different response methods. To determine what effect changing response modalities might have, in **Experiment 4**, we had human participants (n = 29, 1 session each) perform the task using either a touchscreen interface (14,015 total trials; similar to the mice) and an eye-tracking interface (14,485 total trials; similar to the monkeys). To ensure sufficient power, we collected enough participants to detect an effect size of 0.6—less than a quarter of the magnitude of the species-level effect on the probability of switching (Cohen’s d = 2.54, between mice and monkeys) and exploration (Cohen’s d = 2.42, between mice and monkeys). We found no significant differences within participants across response modalities. This was true in performance over chance (**Figure 4D;** paired t-test: t(28) = 0.960, p = 0.35), probability of switching (**Figure 4E**; t(28) = 0.730, p = 0.47), and probability of exploring (**Figure 4F**; t(24) = -0.298, p = 0.77). Thus, the effects of changing response modality are considerably smaller than the differences observed across species and are unlikely to be a sufficient explanation for them.

### Learning index analysis and reward sensitivity

The primates’ tendency to exploit more than mice did not appear to be an artifact of differences in task design or timing. Therefore, we next looked at whether the differences between species were due to their capacity to learn from rewards. We used a “learning index,” a one-trial-back measure normalized by the probability of switching, to measure how rewards affected decisions to switch using data from Experiment 1 (see **Methods**). We found that humans appeared to learn faster compared to mice (**Figure 5A**; humans-mice: p < 0.0001, b = -1.43, 95% CI = [-1.86, -1.00], t(504) = -6.57), but did not differ from monkeys (humans-monkeys: p = 0.047, b = -1.06,95% CI = [-2.12, -0.013], t(345) = -1.99). Monkeys and mice also had a marginal difference in learning speed (**Figure 5A**; monkeys-mice: p < 0.005, b = -0.60, 95% CI = [-1.00, -0.21], t(341) = -2.99). Interpreting this learning index is complicated because it is normalized by the overall probability of switching, and primates switched less often. This means that differences in the learning index may reflect differences in persistence rather than learning from rewards.

**Figure 5.**
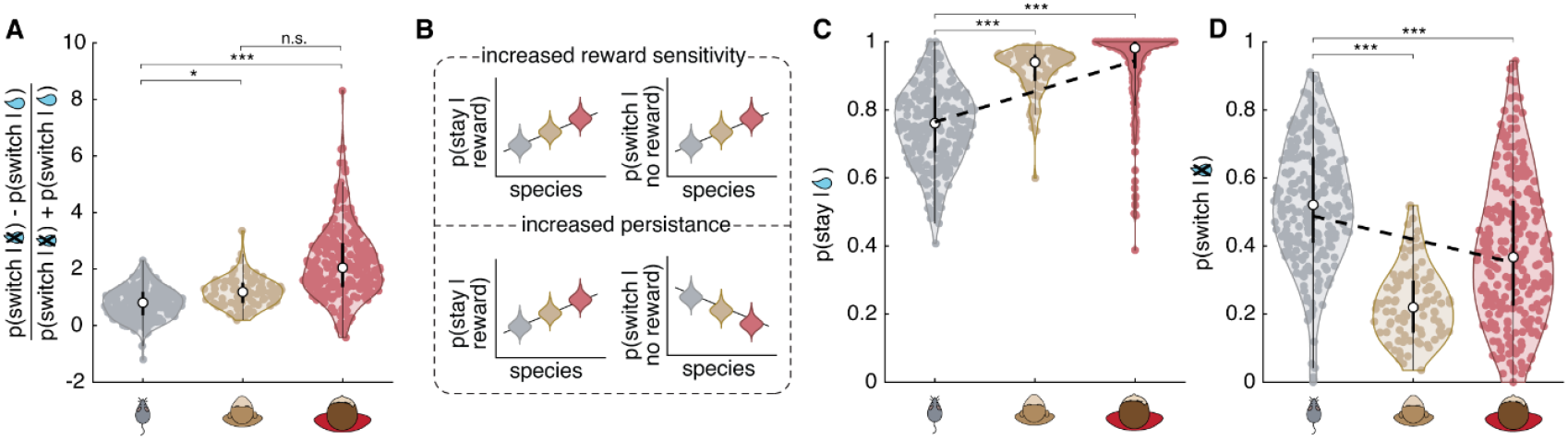
Learning and persistence across species. **A**) Index of reward learning across species. Thick black lines = IQR, thin = whiskers, open circle = median. **B**) Hypothesis cartoon illustrating predictions under the hypothesis that species differences in switching were due to reward sensitivity (top) or persistence (bottom). **C**) Probability of choosing the same option after getting a reward across species. **D**) Probability of choosing a different option after not getting a reward. * Indicates α = 0.05; ** indicates α = 0.001; and *** indicates α = 0.0001.

To differentiate between these explanations, we analysed the choice patterns of rewarded and unrewarded trials. If the difference between species was in learning from rewards, then the tendency to repeat rewarding options should correlate with the tendency to switch away from non-rewarding ones (**Figure 5B**, top). In short, humans should be most sensitive to reward outcomes, followed by monkeys, then mice. Conversely, if the difference between species was in persistence, then the tendency to repeat rewarding options should be inversely related to switching from non-rewarding ones (**Figure 5B**, bottom). In this case, humans would be the most persistent species. Compared to mice, we found that primates persisted more with their choices, both after being rewarded (**Figure 5C**; humans-mice: p < 0.0001, b = -0.18, 95% CI = [-0.21, - 0.15], t(511) = -11,51; monkeys-mice: p < 0.0001, b = -0.20, 95% CI = [-0.28, -0.12], t(344) = - 4.87), and after not receiving a reward (**Figure 5D**; humans-mice: p < 0.0001, b = 0.14, 95% CI = [0.074, 0.20], t(511) = 4.24; monkeys-mice: p < 0.0001, b = 0.32, 95% CI = [0.19, 0.45], t(344) = 4.7). Together, these results suggest the major systematic difference between species was an increase in persistence rather than increased sensitivity to rewards, although some reward sensitivity may also have played a role in humans.

## DISCUSSION

This study compared mice, monkeys, and humans in an uncertain decision-making task. All three species used a similar strategy, alternating between rapidly switching between targets, and persistently choosing the same target. Despite these shared strategies, we found species differences in the average performance and the tendency to switch between targets. Mice switched more frequently than primates. Computational analysis of the switching patterns revealed that the increase in switching in mice was driven by their tendency to exploit less persistently than primates. They initiated exploration more frequently. This was not due to impulsivity nor were species differences a function of low-level differences between the tasks like the number of targets, the timing of the trials, or the modality of the response. Instead, monkeys and humans appeared to persist in exploiting valuable options for longer than mice did.

These results resonate with previous work on differences between primates and rodents. For example, in comparing mice performing a reversal learning task with monkeys performing a blocked bandit task, Woo et al., 2023 found evidence of short- and long-time constants in both species, but also that monkeys underperformed mice, and switched more often. However, these prior results could have been due to species differences or to differences between the tasks. Because the present study harmonised the structure of the task across species, controlled for task differences via additional experiments (i.e., varying the number of targets, trial length, and response modality), and included humans, the results we present here suggest a reinterpretation of that previous study. When the tasks are equated, monkeys outperformed (rather than underperformed) mice, switched less (rather than more), and we did not observe any major differences in reward learning. This could suggest that these specific results in Woo et al., 2023 may have been due to differences in task design. However, both studies agree that the two species share a two-strategy approach to decision-making, even as persistence differs. In our data, persistence, which our computational approach links to exploitative states, is greater in primates compared to mice.

One reason why primates might persist more in exploitation, could be that they have more cognitive self-control: the ability to regulate their impulses, letting them weigh long-term benefits against immediate rewards. The capacity for self-control is more prevalent in species with larger brain sizes (MacLean et al., 2014). Here, self-control could help sustain a choice policy in the absence of reward or help animals avoid the temptation to try something new (Stillman et al., 2017). Indeed, primates were less likely to switch from a target immediately after it did not give a reward, compared to mice. Thus, species differences in the capacity for self-control could help explain why both primates persisted for longer than mice did.

A second, complementary explanation for why primates persisted more than mice could be differences in stability of brain states across species. Single neurons and neural populations process information with characteristic “neural timescales” (Zilio et al., 2021). Previous studies have shown that different brain regions have differing timescales, with longer timescales in the prefrontal cortex compared to sensory cortices (Murray et al., 2014; Golesorkhi et al., 2021; Zilio et al., 2021; Hasson et al., 2008). Brain regions with longer neural timescales are better suited for integrating information over longer periods of time, like in working memory, while brain regions with shorter neural timescales are better suited for processing information that needs quick integration, like sensory cues (Zilio et al., 2021). The prefrontal cortex (PFC) not only has the longest timescales (Murray et al., 2014), but it is also more elaborated in primates compared to mice (Laubach et al., 2018; Preuss and Wise, 2022). The elaborated primate PFC could improve persistence by stabilizing information or control policies in the brain (Cohen et al., 2013; Krawczyk, 2002; Domenech and Koechlin, 2015). This explanation is complimentary because species differences in neural timescales could be the mechanism for species differences in self-control. Future studies are needed to determine how individual differences in self-control and neural timescales predict differences in persistence.

We can also consider a third, ecological reason why primates might persist more than mice: differences in their ecological niches. Social primates, like humans and rhesus macaques, benefit from collective vigilance within their groups (Iki and Kutsukake, 2021), this allows each individual monkey to focus on exploiting resources for longer before looking up to scan for threats. Mice, on the other hand, are mostly prey species (Dickman, 1992) which might require them to be more vigilant and favour less sustained focus on other tasks. Differences in ecological niches across species could imply that we would obtain different results in other environments. Perhaps mice are better adapted to more volatile environments, for example, and would perform better than primates here. The differences in persistence might also be reduced if the task environment were more volatile.

Comparative work is essential both for understanding how the human brain evolved and for ensuring that preclinical research translates into real-world impact. That said, comparative studies also have unique challenges and limitations. For one, whenever data is collected across multiple labs over multiple years, it introduces variability. Here, we worked to harmonise data collection across labs, but it is impossible to equate factors like training time and handling protocols. There were also differences in response and reward modalities between species, due to differences in physicality and experimental apparatuses. While we were able to rule out many factors as sufficient explanations for our major results, we cannot rule out the possibility that task or training differences interacted with real species differences in complex ways. While our results are intriguing, future comparative work will always be needed. In particular, it would be valuable to expand the focus on a single stochastic decision-making task. It would be valuable to know if there are species differences in persistence in other tasks.

Persistence is altered in multiple clinical disorders and by multiple clinically relevant stressors (Verdejo-García et al., 2006; Tolin et al., 2009; Mäntylä et al., 2012; Blanco et al., 2013; Teng et al., 2016; Kaske et al., 2023). This task can be used in both humans and animal models and provides excellent measures of persistence, which is the primary drivers of individual differences in this task (Abbaszadeh et al., 2025). The similarities between species we report here highlight the validity of this general approach for preclinical and translational research. This is especially important because preclinical studies in mice do not always translate well into effective human interventions (Worp et al., 2010; Perrin, 2014; Walker and Eggel, 2020). Despite huge investments in preclinical work on depression in mice, limitations in the behaviours typically assessed in mice as putatively linked with depression have been an impediment to the development of effective and novel treatments (National Institute of Mental Health, 2019; Planchez et al., 2019; Shemesh and Chen, 2023). While our results clearly show that monkeys are a valuable animal model for human conditions in which persistence is compromised, they also introduce a suite of computational tools that can be used to measure persistence in mice—even though that persistence is not as developed. These results further suggest that designing mouse tasks to maximize persistence may go a long way towards improving the relevance and impact of high throughput rodent preclinical testing.

## METHOD

### Experimental models and participant details

All animal care and experimental procedures were approved by the relevant ethical review board (**mice**: the guidelines of the National Institution of Health and the University of Minnesota; **monkeys**: the guidelines of Stanford University Institutional Animal Care and Use Committee and the Rochester University Committee on Animal Resources; **humans for Experiment 1, 3 and 4**: the guidelines of the Comité d’Éthique de la Recherche en Sciences et Santé (CERSES) of the University of Montreal; **humans for Experiment 2**: the guidelines of Princeton University Institutional Review Board). The human data and much of the monkey data has not been analysed or reported previously. Some sessions from two of the five monkeys have been analysed previously (28/58 sessions; (Ebitz et al., 2018). The mouse data has been reported previously (Chen et al., 2021b) but all analyses here are new.

Specific details of each experimental setup are as follows:

#### Mice

Thirty-two BL6129SF1/J mice (16 males and 16 females) were obtained from Jackson Laboratories (stock #101043). Mice arrived at the lab at 7 weeks of age and were housed in groups of four with *ad libitum* access to water and mild food restriction (85–95% of free feeding weight) for the experiment. Animals engaging in operant testing were housed in a 9AM to 9PM reversed light cycle to permit testing during the dark period. Before operant chamber training, animals were food restricted to 85–90% of free feeding body weight. Operant testing occurred five days a week (Monday-Friday). Additional information regarding mouse data collection has been reported previously (Chen et al., 2021b).

#### Monkeys

Five male rhesus macaques (between 5 and 15 years of age; between 6 and 16 kg) participated in this study. Three of the monkeys were singly housed and two were pair housed. All were housed in small colony rooms (6-10 animals per room). Animals were surgically prepared with head restraint prostheses before training began. Analgesics were used to minimise discomfort. After recovery, monkeys were acclimated to the laboratory and head restraint, then placed on controlled access to fluids and trained to perform the task over the course of 3 months. One animal was naive at the start of the experiment, the other four had previously participated in oculomotor and visual attention studies (2 monkeys) or decision-making studies (2 monkeys). Data was collected 5 days a week (Monday-Friday). Additional information regarding two of the monkeys has been reported previously (Ebitz et al., 2018) Data from the other 3 have not been previously analysed or reported.

#### Human

In **Experiment 1, 2, and 3**, humans were recruited via the online platform, Amazon Mechanical (mTurk). To avoid bots and improve data quality, participants were only accepted when they had a minimum of 5000 approved human intelligence tasks (HIT) and a 98% approval rating. Data was collected during specific daytime hours (9 AM to 2 PM EST for **Experiments 1 and 3**, and 9 AM to 5 PM EST for **Experiments 2 and 4**). Participants received a base payment of $0.50 USD plus a performance-based bonus of $0.02 per rewarded trial. The total compensation was $4.35 mean ± 0.90 USD (mean ± SD) for all 3 online experiments (n = 707), with payments ranging from a minimum of $1.54 USD to a maximum of $9.22 USD. In **Experiment 1**, participants self-reported as 120 females, 137 males, and 1 preferred not to say. For **Experiment 2**, 150 participants completed the task (gender not collected). In **Experiment 3** participants self-reported as 139 females, 158 males, and 2 preferred not to say. In Experiment 4, participants completed the task in-person at the lab and were compensated $12 CAD per hour, plus a $0.02 CAD bonus for each rewarded trial. The total compensation was $31.86 mean ± 3.22 CAD (mean ± SD) for the in-person experiment, with payments ranging from a minimum of $26.3 CAD to a maximum of $39.94 CAD. A total of 29 participants completed the task. Data was collected from 9AM to 5PM EST. For each participant, the first response modality in which they performed—eye tracking versus touchscreen—was randomized.

#### Procedure

For each experiment, participants performed a spatial restless k-armed bandit task where they chose between physically identical visual cues. In this task, physically identical targets are presented in spatial locations that are associated with a probability of reward. Reward probabilities ranged between 0.9 and 0.1 and could diminish or increase over time at a rate that was fixed across experiments (10% chance of a step of 0.1). For 2/5 monkeys, the floor probability of reward was 0.3, rather than 0.1, to improve motivation (see **Results**).

Rewards were variable, independent, and probabilistic. Participants could only infer values through sampling the targets and integrating reward history over multiple trials. There were minor variations between the mouse, monkey, and human studies due to a combination of factors: (1) the data was collected independently across multiple labs, (2) the tasks were adapted to the typical research approaches used in each species. Mice and monkeys both received a primary, liquid reinforcer as reward. Humans received money, a secondary reinforcer. Monkeys received a 3-target version of the bandit task, mice received a 2-target version, and humans received a 2-, 3- and 4-target version. Additional variations between the tasks are described below:

#### Mice

Mice indicated their choices by nose-poking on a touchscreen. Each trial, mice were presented with a cue displaying two identical squares positioned on the left and right side of the screen. A nose poke to one of the target locations on the touchscreen was required to register a response. Rewards were given in the form of food pellets. Mice completed either 300 trials or spent a maximum of two hours in the operant chamber. On average, mice performed 276.38 trials (min: 46 trials, max: 300 trials) per session.

#### Monkeys

Monkeys indicated their choices by making saccadic eye movements towards one of three identical gabor kernels. Choices were registered when the monkeys fixated on the eccentric target for a specified minimum period (150ms). Eye position was monitored at 1000Hz via an infrared eye tracker (SR Research). Rewards were given in the form of juice. On average, monkeys performed 622.34 trials (min: 144 trials, max: 1377 trials) per session.

#### Humans

Humans indicated their choices by moving a computer mouse towards one of the backs of playing cards on the screen. In Experiment 1, human participants had to choose between 2 identical blue backs of playing cards. In Experiment 2, human participants had to choose between 2, 3, or 4 backs of playing cards, with each card being identical except for their colour. In Experiment 3, human participants had to choose between 3 identical blue backs of playing cards. They used a computer mouse to click the desired options and register their response. Rewards were given in the form of money ($0.02 per reward). Every human participant completed 300 trials per session, except for 1 participant who completed 299 trials during their session. Prior to the experiment, the humans completed an additional 20-25 practise trials, which were meant to familiarise them with the task but were not included in the analyses. In Experiment 4, each participant completed two successive versions of the task. In the touchscreen version, they selected targets by physically touching the display, also touching on a fixation cross located in the middle of the screen during the ITI. In the eye-tracking version, their head position was stabilized with a chin rest while an SR Research infrared eye tracker recorded their eye movements. The order of response modalities was randomized for each participant.

### Quantification and Statistical Analysis

#### General analysis techniques

Data was analysed with custom MATLAB scripts and p-values were compared against the standard α = 0.05 threshold for species-specific paired t-tests, and one-way ANOVAs. When conducting the three planned pairwise species comparisons for Experiment 1 (humans vs. mice, monkeys vs. mice, and humans vs. monkeys), we applied a Bonferroni correction to control the family-wise error rate (α′ = 0.05/3 ≈ 0.017). We analysed the data using a general linear mixed-effects model, with Species as a fixed factor and subjects nested within species as a random factor to account for repeated measures on the same animal. In cases where we compared mice vs monkeys separately, we included sessions as a fixed factor (plus its interaction with species), however, while these were included in the model, we did not report the session and interaction effect in the paper. The Session term wasn’t included for humans, since each human completed only a single session. In **Experiment 1**, the sample size for mice (n = 32; total sessions = 255), for monkeys n = 5, total sessions = 93, for humans (2-armed bandit task) n = 258, total sessions = 258. In **Experiment 2**, the sample size for humans was (2-armed bandit task) n = 50, total sessions = 50, for humans (3-armed bandit task) n = 50, total sessions = 50, for humans (4-armed bandit task) n = 50, total sessions = 50. In **Experiment 3**, the sample size for humans was (ITI 200ms) n = 99, total sessions = 99, for humans (ITI 2000ms) n = 93, total sessions = 93, for humans (ITI 4000ms) n = 107, total sessions = 107. In **Experiment 4**, the sample size for humans was (2-armed touchscreen and eye-tracking bandit task) n = 29, total sessions = 29.

A small number of sessions from some participants were excluded from analysis. Specifically, 6 sessions out of 150 were excluded in **Experiment 2** because participants did not select all available targets during the session and this experiment specifically looked at the behavioural effects of manipulating the number of targets. Otherwise, sessions were only excluded from specific analyses when these analyses were impossible, given the participants’ behaviour. For example, in **Experiment 1**, 7 sessions out of 606 (3 for mice, 0 for monkeys, 4 for humans) were excluded from certain analyses of switching behaviour (i.e. the learning index, mixture model, HMM) because the participant did not switch during those sessions. However, these sessions were included in all other analyses, including of the probability of switching (i.e. **Figure 1D**). The specific exclusion criteria for each analysis as well as the number of excluded sessions is described within the relevant section of the Methods. For response time analyses, any trial exceeding 1.5 times the interquartile range (IQR) was excluded as an outlier. Response time analyses excluded 14 sessions from 93 monkeys because no reaction time data could be recovered from these sessions.

#### Random Exploration Among k-Arms

In a random exploration strategy, a target is selected at random on each trial. This means that the probability of repeating a choice is the same as the independent probability of making that choice (i.e. it is always 1/k, where k is the number of options). The probability of switching away from a previous option is then the probability of choosing any other option:

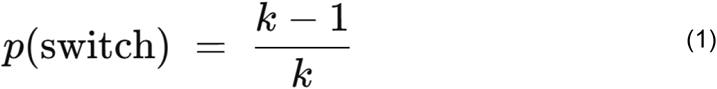

Note that as k increases, as the number of targets increases, the probability of switching increases systematically, under the hypothesis that decisions are made randomly. This is the explicit equation, with 0 free parameters, that is plotted in **Figure 3F**.

#### Analysing the modulated indices

In order to determine if response time would reveal species differences in task strategy, we looked at how response time changed as a function of certain task events. Because species performed slightly different variations of the task, with differing response modalities (i.e. touch screen for mice, saccades for monkeys, and computer mouse for humans), we were unable to compare raw response times directly. Instead, we computed “modulated indices of reaction times” by normalising the averaged difference in response times between 2 conditions by the standard deviation of response time. We specifically assessed whether response time was modulated by exploration (versus exploitation) and by reward (versus no reward) in any species. If response time was faster in exploration trials, as opposed to exploitation trials, in some species, the modulated index would be above 0, implying that exploration is “impulsive” (faster). If response time is slower for exploration, the modulated index would be below 0, implying that exploration is a slower, more “deliberate” choice process relative to exploitation (**Figure 2G** - top). A similar approach was applied to compare rewarded and unrewarded trials (**Figure 2G** - bottom). The explore/exploit modulation index could only be calculated in sessions where (1) response time data was available, and (2) that included both exploration and exploitation (560 total sessions / 606 sessions). Analyses of the rewarded/unrewarded modulation index included all sessions where response time data was available (592 total sessions / 606 sessions).

## REFERENCES

Abbaszadeh M, Ozanick E, Magen N, Darrow D, Yan X, Grissom N, Herman AB, Ebitz B (2025) Individual differences in sequential decision-making. BioRxiv Prepr Serv Biol: 2025.04.04.647306.

Arrhenius S (1889) Über die Reaktionsgeschwindigkeit bei der Inversion von Rohrzucker durch Säuren. Z Für Phys Chem 4U:226–248.

Barger KJ-A (2006) Mixtures of Exponential Distributions to Describe the Distribution of Poisson Means in Estimating the Number of Unobserved Classes. Available at: https://ecommons.cornell.edu/handle/1813/2953 [Accessed June 16, 2022].

Bari BA, Grossman CD, Lubin EE, Rajagopalan AE, Cressy JI, Cohen JY (2019) Stable Representations of Decision Variables for Flexible Behavior. Neuron 103:922-933.e7.

Bilmes J (2000) A Gentle Tutorial of the EM Algorithm and its Application to Parameter Estimation for Gaussian Mixture and Hidden Markov Models. Tech Rep ICSI-TR-97-021 Univ Berkeley 4.

Blanco NJ, Otto AR, Maddox WT, Beevers CG, Love BC (2013) The influence of depression symptoms on exploratory decision-making. Cognition 129:563–568.

Boltzmann L (1868) Studien uber das gleichgewicht der lebenden kraft. Wissenschafiliche Abh 1:49–96.

Boyden ES, Zhang F, Bamberg E, Nagel G, Deisseroth K (2005) Millisecond-timescale, genetically targeted optical control of neural activity. Nat Neurosci 8:1263–1268.

Burnham KP, Anderson DR (2002) Model selection and multimodel inference: a practical information-theoretic approach, 2nd ed. New York: Springer.

Chen CS, Ebitz RB, Bindas SR, Redish AD, Hayden BY, Grissom NM (2021a) Divergent Strategies for Learning in Males and Females. Curr Biol CB 31:39-50.e4.

Chen CS, Knep E, Han A, Ebitz RB, Grissom NM (2021b) Sex differences in learning from exploration Izquierdo A, Wassum KM, Izquierdo A, eds. eLife 10:e69748.

Cohen JR, Berkman ET, Lieberman MD (2013) Intentional and Incidental Self-Control in Ventrolateral Prefrontal Cortex. In: Principles of Frontal Lobe Function (Tranel D, Stuss DT, Knight RT, eds), pp 0. Oxford University Press. Available at: 10.1093/med/9780199837755.003.0030 [Accessed April 19, 2024].

Dickman CR (1992) Predation and Habitat Shift in the House Mouse, Mus Domesticus. Ecology 73:313–322.

Disotell TR, Tosi AJ (2007) The monkey’s perspective. Genome Biol 8:226.

Domenech P, Koechlin E (2015) Executive control and decision-making in the prefrontal cortex. Curr Opin Behav Sci 1:101–106.

Ebitz RB, Albarran E, Moore T (2018) Exploration Disrupts Choice-Predictive Signals and Alters Dynamics in Prefrontal Cortex. Neuron 97:450-461.e9.

Ebitz RB, Sleezer BJ, Jedema HP, Bradberry CW, Hayden BY (2019) Tonic exploration governs both flexibility and lapses. PLoS Comput Biol 15:e1007475.

Ebitz RB, Tu JC, Hayden BY (2020) Rules warp feature encoding in decision-making circuits. PLOS Biol 18:e3000951.

Ellenbroek B, Youn J (2016) Rodent models in neuroscience research: is it a rat race? Dis Model Mech 9:1079–1087.

Ernst PB, Carvunis A-R (2018) Of mice, men and immunity: a case for evolutionary systems biology. Nat Immunol 19:421–425.

Gibbs RA et al. (2007) Evolutionary and biomedical insights from the rhesus macaque genome. Science 316:222–234.

Golesorkhi M, Gomez-Pilar J, Zilio F, Berberian N, Wolff A, Yagoub MCE, Northoff G (2021) The brain and its time: intrinsic neural timescales are key for input processing. Commun Biol 4:1–16.

Grabenhorst F, Tsutsui K-I, Kobayashi S, Schultz W (2019) Primate prefrontal neurons signal economic risk derived from the statistics of recent reward experience Lee D, Ivry RB, Louie K, eds. eLife 8:e44838.

Groman SM, Smith NJ, Petrullli JR, Massi B, Chen L, Ropchan J, Huang Y, Lee D, Morris ED, Taylor JR (2016) Dopamine D3 Receptor Availability Is Associated with Inflexible Decision Making. J Neurosci Off J Soc Neurosci 36:6732–6741.

Grossman CD, Bari BA, Cohen JY (2022) Serotonin neurons modulate learning rate through uncertainty. Curr Biol 32:586-599.e7.

Hasson U, Yang E, Vallines I, Heeger DJ, Rubin N (2008) A Hierarchy of Temporal Receptive Windows in Human Cortex. J Neurosci 28:2539–2550.

Iki S, Kutsukake N (2021) Japanese macaques relax vigilance when surrounded by kin. Anim Behav 179:173–181.

Iyer ES, Weinberg A, Bagot RC (2022) Ambiguity and conflict: Dissecting uncertainty in decision-making. Behav Neurosci 136:1–12.

Izquierdo A, Aguirre C, Hart EE, Stolyarova A (2019) Rodent Models of Adaptive Value Learning and Decision-Making. Methods Mol Biol Clifton NJ 2011:105–119.

Kaske EA, Chen CS, Meyer C, Yang F, Ebitz B, Grissom N, Kapoor A, Darrow DP, Herman AB (2023) Prolonged physiological stress is associated with a lower rate of exploratory learning that is compounded by depression. Biol Psychiatry Cogn Neurosci Neuroimaging 8:703–711.

Knox W, Otto A, Stone P, Love B (2012) The Nature of Belief-Directed Exploratory Choice in Human Decision-Making. Front Psychol 2 Available at: https://www.frontiersin.org/articles/10.3389/fpsyg.2011.00398 [Accessed October 24, 2023].

Krawczyk DC (2002) Contributions of the prefrontal cortex to the neural basis of human decision making. Neurosci Biobehav Rev 26:631–664.

Lau B, Glimcher PW (2005) Dynamic Response-by-Response Models of Matching Behavior in Rhesus Monkeys. J Exp Anal Behav 84:555–579.

Laubach M, Amarante LM, Swanson K, White SR (2018) What, If Anything, Is Rodent Prefrontal Cortex? eNeuro 5 Available at: https://www.eneuro.org/content/5/5/ENEURO.0315-18.2018 [Accessed April 19, 2024].

MacLean EL et al. (2014) The evolution of self-control. Proc Natl Acad Sci 111:E2140–E2148.

Manger P, Cort J, Ebrahim N, Goodman A, Henning J, Karolia M, Rodrigues S-L, Strkalj G (2008) Is 21st century neuroscience too focussed on the rat/mouse model of brain function and dysfunction? Front Neuroanat 2 Available at: https://www.frontiersin.org/articles/10.3389/neuro.05.005.2008 [Accessed October 18, 2023].

Mäntylä T, Still J, Gullberg S, Del Missier F (2012) Decision Making in Adults With ADHD. J Atten Disord 16:164–173.

Mehlhorn K, Newell BR, Todd PM, Lee MD, Morgan K, Braithwaite VA, Hausmann D, Fiedler K, Gonzalez C (2015) Unpacking the exploration–exploitation tradeoff: A synthesis of human and animal literatures. Decision 2:191–215.

Murray JD, Bernacchia A, Freedman DJ, Romo R, Wallis JD, Cai X, Padoa-Schioppa C, Pasternak T, Seo H, Lee D, Wang X-J (2014) A hierarchy of intrinsic timescales across primate cortex. Nat Neurosci 17:1661–1663.

National Institute of Mental Health (2019) NOT-MH-19-053: Notice of NIMHs Considerations Regarding the Use of Animal Neurobehavioral Approaches in Basic and Pre-clinical Studies. Available at: https://grants.nih.gov/grants/guide/notice-files/NOT-MH-19-053.html [Accessed May 26, 2025].

Perrin S (2014) Preclinical research: Make mouse studies work. Nature 507:423–425.

Planchez B, Surget A, Belzung C (2019) Animal models of major depression: drawbacks and challenges. J Neural Transm Vienna Austria 1996 126:1383–1408.

Preuss TM, Wise SP (2022) Evolution of prefrontal cortex. Neuropsychopharmacol Off Publ Am Coll Neuropsychopharmacol 47:3–19.

Rich AS, Gureckis TM (2018) Exploratory choice reflects the future value of information. Decision 5:177–192.

Saddoris MP, Sugam JA, Stuber GD, Witten IB, Deisseroth K, Carelli RM (2015) Mesolimbic Dopamine Dynamically Tracks, and Is Causally Linked to, Discrete Aspects of Value-Based Decision Making. Biol Psychiatry 77:903–911.

Shemesh Y, Chen A (2023) A paradigm shift in translational psychiatry through rodent neuroethology. Mol Psychiatry 28:993–1003.

Shourkeshti A, Marrocco G, Jurewicz K, Moore T, Ebitz RB (2023) Pupil size predicts the onset of exploration in brain and behavior. BioRxiv Prepr Serv Biol:2023.05.24.541981.

Siegal ML, Bergman A (2002) Waddington’s canalization revisited: Developmental stability and evolution. Proc Natl Acad Sci 99:10528–10532.

Soltani A, Izquierdo A (2019) Adaptive learning under expected and unexpected uncertainty. Nat Rev Neurosci 20:635–644.

Stevenson TJ, Alward BA, Ebling FJP, Fernald RD, Kelly A, Ophir AG (2018) The Value of Comparative Animal Research: Krogh’s Principle Facilitates Scientific Discoveries. Policy Insights Behav Brain Sci 5:118.

Stillman PE, Medvedev D, Ferguson MJ (2017) Resisting Temptation: Tracking How Self-Control Conflicts Are Successfully Resolved in Real Time. Psychol Sci 28:1240–1258.

Teng C, Otero M, Geraci M, Blair RJR, Pine DS, Grillon C, Blair KS (2016) Abnormal decision-making in generalized anxiety disorder: Aversion of risk or stimulus-reinforcement impairment? Psychiatry Res 237:351–356.

Tolin DF, Kiehl KA, Worhunsky P, Book GA, Maltby N (2009) An exploratory study of the neural mechanisms of decision making in compulsive hoarding. Psychol Med 39:325–336.

Trepka E, Spitmaan M, Bari BA, Costa VD, Cohen JY, Soltani A (2021) Entropy-based metrics for predicting choice behavior based on local response to reward. Nat Commun 12:6567.

Verdejo-García A, Pérez-García M, Bechara A (2006) Emotion, Decision-Making and Substance Dependence: A Somatic-Marker Model of Addiction. Curr Neuropharmacol 4:17–31.

Waddington CH (1942) Canalization of Development and the Inheritance of Acquired Characters. Nature 150:563–565.

Walker RL, Eggel M (2020) From Mice to Monkeys? Beyond Orthodox Approaches to the Ethics of Animal Model Choice. Animals 10:77.

Wilson RC, Bonawitz E, Costa VD, Ebitz RB (2021) Balancing exploration and exploitation with information and randomization. Curr Opin Behav Sci 38:49–56.

Woo JH, Aguirre CG, Bari BA, Tsutsui K-I, Grabenhorst F, Cohen JY, Schultz W, Izquierdo A, Soltani A (2023) Mechanisms of adjustments to different types of uncertainty in the reward environment across mice and monkeys. Cogn Affect Behav Neurosci 23:600–619.

Worp HB van der, Howells DW, Sena ES, Porritt MJ, Rewell S, O’Collins V, Macleod MR (2010) Can Animal Models of Disease Reliably Inform Human Studies? PLOS Med 7:e1000245.

Yan X, Ebitz RB, Grissom N, Darrow DP, Herman AB (2023) A low dimensional manifold of human exploratory behavior reveals opposing roles for apathy and anxiety. BioRxiv Prepr Serv Biol:2023.06.19.545645.

Zilio F, Gomez-Pilar J, Cao S, Zhang J, Zang D, Qi Z, Tan J, Hiromi T, Wu X, Fogel S, Huang Z, Hohmann MR, Fomina T, Synofzik M, Grosse-Wentrup M, Owen AM, Northoff G (2021) Are intrinsic neural timescales related to sensory processing? Evidence from abnormal behavioral states. NeuroImage 226:117579.

