## Supplemental methods, figures, and tables for "Persistent Decision-Making in Mice, Monkeys, and Humans"

*Exponential Mixture Distribution*

We analysed the temporal structure of the participants' choice sequences with a mixture model. If a single time constant (probability of switching) governed the behaviour, we would expect to see exponentially distributed inter-switch intervals. That is, the distribution of inter-switch intervals should be well described by the following model:

$$f(x) = \frac{1}{\beta} e^{-\frac{x}{\beta}} \quad (1)$$

Where β is the "survival parameter" of the model: the average inter-switch interval. However, although the time between switch decisions was largely monotonically decreasing and concave upwards, the distribution was not well described by a single exponential distribution (**Figure 2A**). Participants had more short-latency and more long latency choice runs, indicating that a single switching probability could not have generated the data. Therefore, we fit mixtures of varying numbers of exponential distributions (1-4) to all species (**Figure 2A**), in order to infer the number of switching regimes in these choice processes. For continuous-time processes, these mixture distributions would be of the form:

$$f(x) = \sum_{i=1}^{n} \pi_i e^{-\frac{x}{\beta_i}} \quad (2)$$

Where $1 \geq \pi_i \geq 0$ for all $\pi_i$, and $\Sigma_i \pi i = 1$. Here, each $\beta_i$ reflects the survival parameter (average inter-switch interval) for each component distribution i and the $\pi_i$ reflects the relative weight of each component. Because trials were discrete, we fit the discrete analog of this distribution: mixtures of 1-4 discrete exponential (geometric) distributions (Barger, 2006). Mixtures were fit via the expectation-maximisation algorithm and we used standard model comparison (Burnham and Anderson, 2002) to determine the most probable number of mixing components (**Figure 2A, Results**).

We used a bootstrap procedure to illustrate the distribution of mixture model parameters in **Figure 2B** and **Figure 3F**. This meant that we resampled, with replacement, from the sessions collected in each species to generate bootstrapped distributions of run lengths (n distributions = 1000, number of sessions equal to the data). We then fit the exponential mixture model to each sample of run lengths, giving a bootstrapped estimate of mixture model parameters. (N.B. Statistical analyses were done on the raw, non-bootstrapped, data, the bootstrapping was only done for illustration.)

Some participants had to be excluded from mixture model analyses because their distribution of run lengths prevented the identification of model parameters. This could happen either because they either had fewer than 2 switches between options (i.e. it was impossible to measure any run lengths) or because their run lengths lacked the variation required for the expectation maximisation algorithm to function (i.e. all run lengths were identical). In **Experiment 1**, 4 human sessions (of 606) were excluded (11 total, when including the 7 excluded previously because no switches were observed) in the main results. In **Figure S2B** of the supplementary materials, sessions were divided into their first 150 and last 150 trials, and any session with no switches in either half was excluded—removing an additional 6 mouse, 6 monkey, and 39 human sessions. In **Experiment 2**, 2 sessions out of 150 were excluded (8 total, including the 6 excluded previously for not choosing all available targets).

*Hidden Markov Model (HMM)*

In order to identify how often different species were exploring or exploiting, we fit an HMM to each session of each species. Here, choices (y) are "emissions" that are generated by an unobserved decision process that is in some latent, hidden state (z). Latent states are defined by both the probability of making each choice k (out of $N_k$ possible options), and by the probability of transitioning from each state to every other state. Our model consisted of two types of states, the explore state and the exploit state. The emissions model for the explore state was uniform across the options:

$$p(y_t = k | z_t = explore) = \frac{1}{N_k} \quad (3)$$

This is the maximum entropy distribution for a categorical variable—the distribution that makes the fewest number of assumptions about the true distribution and thus does not bias the model towards or away from any particular type of high-entropy choice period. This does not require, imply, impose, or exclude that decision-making happening under exploration is random (Ebitz et al., 2019, 2020). Because exploitation involves repeated sampling of each option, exploit states only permitted choice emissions that matched one option. That is:

$$\begin{cases} p(y_t = k | z_t = exploit_i, k \in exploit_i) = 1 \\ p(y_t = k | z_t = exploit_i, k \notin exploit_i) = 0 \end{cases} \quad (4)$$

The latent states in this model are Markovian, meaning that they are time-independent. They depend only on the most recent state ($z_t$):

$$p(z_t | z_{t-1}, y_{t-1}, \ldots, z_1, y_1) = p(z_t | z_{t-1}) \quad (5)$$

This means that we can describe the entire pattern of dynamics in terms of a single transition matrix. This matrix is a system of stochastic equations describing the one-time- step probability of transitioning between every combination of past and future states (i, j).

$$p(z_t = i | z_{t-1} = j) \quad (6)$$

Because the task varied in the number of available targets, the number of states scaled accordingly. In the version with two targets, participants could occupy three states (one explore state and two exploit states; i.e. one for each target). In the three-target version, they had four states (one explore state and three exploit states). And in the four-target version, they had five states in total (one explore state alongside four exploit states). To produce long, exponentially-distributed runs of repeated choices to a single target, the HMM had one latent exploitative state for each target. To produce short, random run lengths, the HMM had one shared explore state from which decisions to any of the choices were equally likely. For all participants, parameters were tied across exploit states such that each exploit state had the same probability of beginning (from exploring) and of sustaining itself. Transitions out of the exploration, into exploitative states, were similarly tied. The model also assumed that the participants had to pass through exploration in order to start exploiting a new option, even if only for a single trial. This is because the utility of exploration is to maximise information about the environment (Mehlhorn et al., 2015). If a participant switches from a bout of exploiting one option to another option, that very first trial after switching should be exploratory because the outcome or reward contingency of that new option is unknown and that behaviour of switching aims to gain information. Through fixing the emissions model, constraining the structure of the transition matrix, and tying the parameters, the final HMM had only two free parameters: one corresponding to the probability of exploring, given exploration on the last trial, and one corresponding to the probability of exploiting, given exploitation on the last trial.

The model was fit via expectation-maximisation using the Baum Welch algorithm (Bilmes, 2000). This algorithm finds a (possibly local) maxima of the complete-data likelihood. A complete set of parameters θ includes the emission and transition models, discussed already, but also initial distribution over states. Because the participants had no knowledge of the environment at the first trial of the session, we assumed they began by exploring, rather than adding another parameter to the model here. The algorithm was reinitialized with random seeds 20 times, and the model that maximised the observed (incomplete) data log likelihood across all the sessions for each animal was ultimately taken as the best. To ultimately infer latent states from choices, we used the forward-backward algorithm to label the most probable sequence of latent states.

Some participants were excluded from analyses that depended on the HMM because the model did not fit these participants. This totalled 59 sessions out of 1084 (>5.5%, 17 for mice, 0 for monkeys, 15 for humans in **Experiment 1**, 9 in **Experiment 2**, 14 in **Experiment 3**, and 4 in **Experiment 4**). The HMM model could fail to fit for 2 reasons: (1) because participants only chose a single target for the whole session (making model parameters unidentifiable) or (2) because fitting procedure resulted in a solution that violated the assumption of longer choice runs under exploitation compared to exploration (where the probability of stopping a bout of exploitation was an obvious outlier in the distribution of this parameter across all species; threshold for exclusion set at 0.4).

*Analysing HMM Dynamics (State Dynamic Landscapes)*

In order to understand the dynamics of exploration and exploitation, we analysed the HMMs. Here, we use the term "dynamics" to mean the equations that govern how a system evolves over time. In fitting our HMMs, we were fitting a set of equations that describe these dynamics: the probability of transitions between exploration and exploitation and vice versa. To illustrate how goal dynamics differed across groups, we performed certain thermodynamic analyses of the long-term behaviour of the fitted equations, generating insight into the potential energy of each state in each species (**Figure 2C**).

In statistical mechanics, processes within a system (like a decision-maker at some moment in time) occupy states (like exploration or exploitation). States have energy associated with them, related to the long-time scale probability of observing a process in those states. A low-energy state is one that is very stable and deep, much like a valley between two mountain peaks. Low-energy states will be over-represented in the system's long-term behaviour. A high energy state, like the top of a mountain, is less stable. High-energy states will be under-represented in the system's behaviour. The probability of observing a process in a given state i is related to the energy of that state ($E_i$) via the Boltzman distribution:

$$p_i = \frac{1}{Z} e^{\frac{-E_i}{k_B T}} \tag{7}$$

where Z is the partition function of the system, $k_B$ is the Boltzman constant, and T is the temperature. If we focus on the ratio between two state probabilities, the partition functions cancel out and the relative occupancy of the two states is now a function of the difference in energy between them:

$$\frac{p_i}{p_j} = e^{\frac{-\left(E_i - E_j\right)}{k_B t}} \tag{8}$$

Rearranging, we express the difference in energy between two states as a function of the difference in the long-term probability of those states being occupied:

$$ln\left(\frac{p_i}{p_j}\right) k_B T = E_j - E_i \tag{9}$$

Meaning that the difference in the energetic depth of the states (the Gibbs Free Energy) is proportional to the natural log of the probability of each state, up to some multiplicative factor $k_B T$. To calculate the probability of exploration and exploitation ($p_i$ and $p_j$), we solved for the stationary distribution of the fitted HMMs. The stationary distribution is the equilibrium probability distribution over states. This means that this distribution is the relative frequency of each state that we would observe if the model's dynamics were run for an infinite period of time. Each entry of the model's transition matrix reflects the probability that the participant would move from one state (e.g. exploring) to another (e.g. exploiting) at each moment in time. Because the parameters for all the exploitation states were tied, each transition matrix effectively had two states—an explore state and a generic exploit that described the dynamics of all exploit states. Each of the k sessions had its own transition matrix ($A_k$), which describes how the entire system—an entire probability distribution over states—would evolve from time point to time point. We observe how the dynamics evolve any probability distribution over states (π) by applying the dynamics to this distribution:

$$\pi_{t+1} = \pi_t A_k \quad (10)$$

Over many time steps, ergodic systems reach a point where the state distributions are unchanged by continued application of the transition matrix as the distribution of states reaches its equilibrium. That is, in stationary systems, there exists a stationary distribution, π, such that:

$$\pi = \pi A_k \quad (11)$$

If it exists, this distribution is a (normalised) left eigenvector of the transition matrix $A_k$ with an eigenvalue of 1, so we solved for this eigenvector to determine the stationary distribution of each $A_k$. We then took an average of these stationary distributions across all sessions for each group, and plugged these back into the Boltzman equations to calculate the relative energy (depth) of exploration and exploitation as illustrated in **Figure 2F** and **Figure 3D**.

In order to understand the dynamics of exploration and exploitation, we need to not only understand the depth of the two states, but also the height of the energetic barrier between them: the energy required to transition from exploration to exploitation and back again. Here, we build on the Arrhenius equation from chemical kinetics that relates the rate of transitions (k) between some pair of states to the activation energy required to affect these transitions ($E_a$):

$$k = Ae^{\frac{E_a}{k_B T}} \quad (12)$$

where A is a constant pre-exponential factor related to the readiness of reactants to undergo the transformation. We set this to one. Again, $k_B T$ is the product of temperature and the Boltzman constant. Note the similarities between this equation and the Boltzman distribution illustrated earlier. Rearranging to solve for activation energy yields:

$$E_a = -ln\left(\frac{k}{A}\right) k_B T \quad (13)$$

Thus, activation energy, much like the relative depth of each state, is also proportional to some measurable function of behaviour, up to some multiplicative factor $k_BT$. Note that our approach has only identified the energy of three discrete states (an explore state, an exploit state, and the peak of the barrier between them). These are illustrated by tracing a continuous potential through these three points to provide a physical intuition for the differences in explore/exploit dynamics between species.

To create the attractor basin graphs, transition matrices were calculated individually for all participants (Seed = 20), and then averaged across groups, see Methods section: Analysing HMM Dynamics (state dynamic landscapes) for more details. All statistical tests used and statistical details were reported in the results.

### Supplementary Figures and Tables

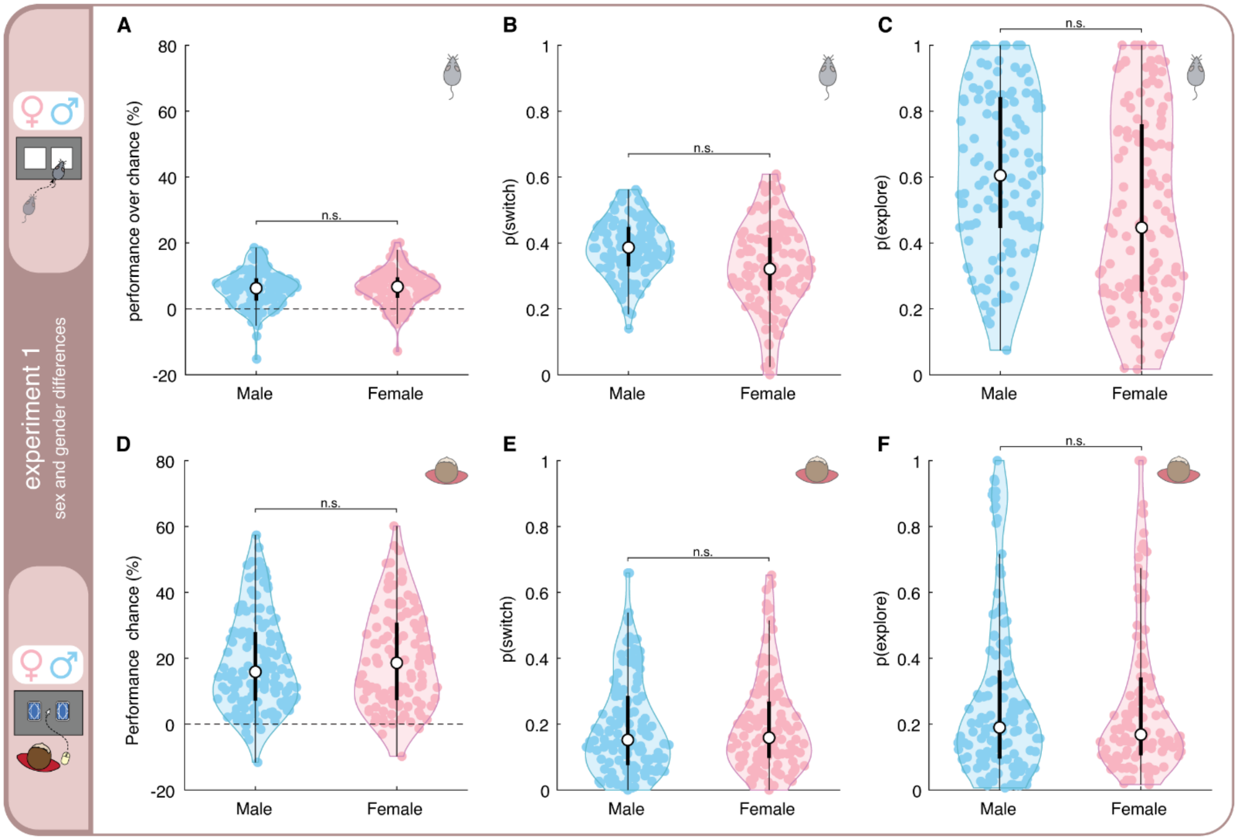


**Figure S1. No sex and gender differences found across participants. A)** Percentage of reward relative to chance across self-reported genders in humans. Thick black lines = IQR, thin = whiskers, open circle = median. **B**) Switch probability by gender. **C**) Probability of exploration by gender. **D-E-F**) Same as in A-B-C) but for mice. Asterisks represent significance levels as follows: * indicates standard $\alpha = 0.05$ threshold; ** indicates $\alpha = 0.001$; and *** indicates $\alpha = 0.0001$.

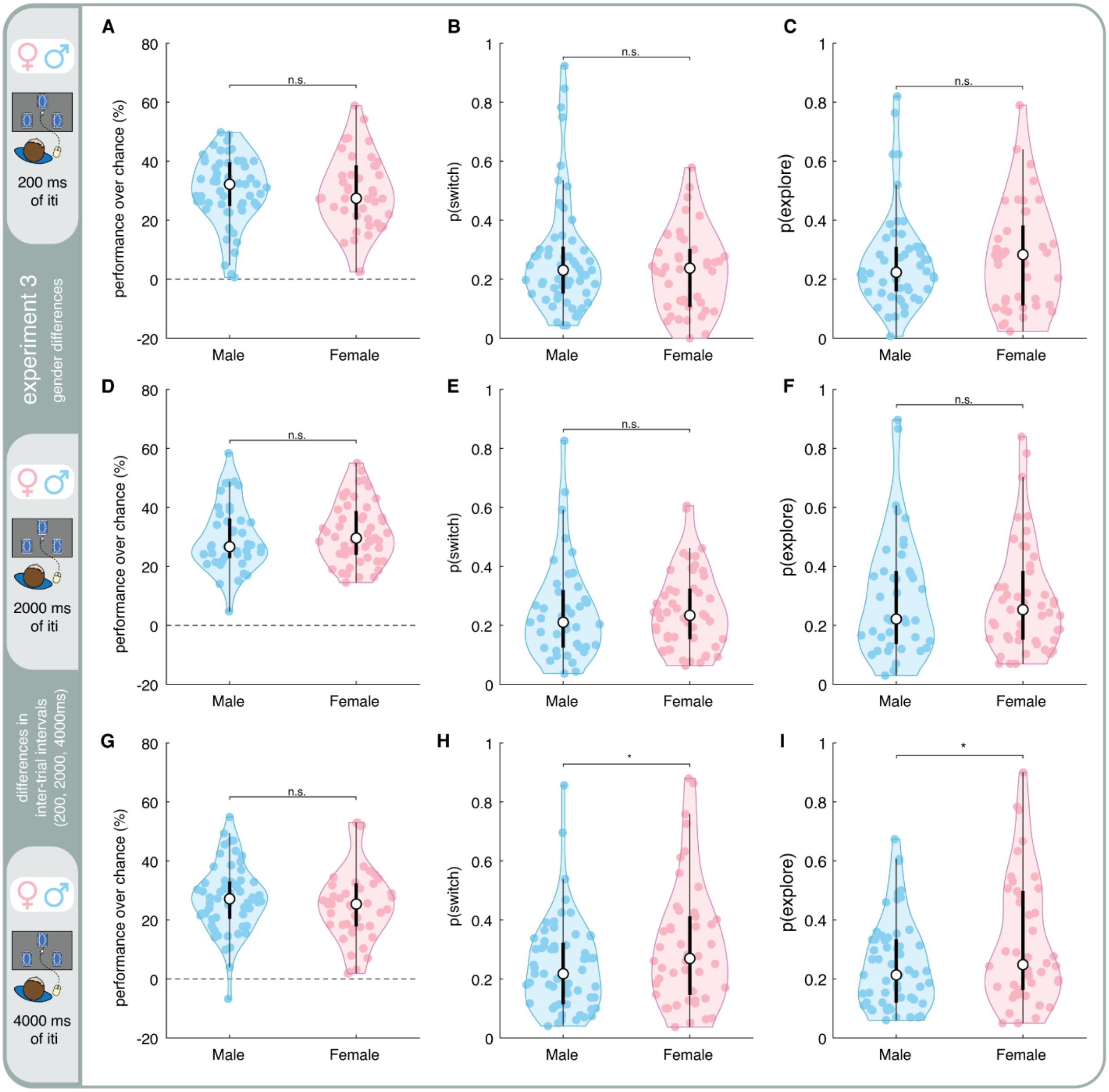


**Figure S2. Sex differences across participants. A**) Percentage of reward relative to chance across self-reported genders in humans doing a 3-armed bandit task with an ITI of 200ms **B**) Switch probability by gender in humans doing a 3-armed bandit task with an ITI of 200ms. **C**) Probability of exploration by gender in humans doing a 3-armed bandit task with an ITI of 200ms. **D-E-F)** Same as in **A-B-C)** but for humans performing a bandit task with an ITI of 2000ms. **G-H-I)** Same as in **A-F)** but for humans performing a bandit task with an ITI of 4000ms. Thick black lines = IQR, thin = whiskers, open circle = median. Asterisks represent significance levels as follows: * indicates standard α = 0.05 threshold; ** indicates α = 0.001; and *** indicates α = 0.0001.

| | 1 component, 1 parameter mixture model | 2 component, 3 parameter mixture model | likelihood ratio test |
|---|---|---|---|
| Mice | –43,757.9 | -40,960.5 | LRT $\chi^2(2)=5594.78$; $p < 0.0001$ |
| Monkeys | -23,253.0 | -19,561.9 | LRT $\chi^2(2)=7382.16$; $p < 0.0001$ |
| Humans | -34,644.8 | -31,048.3 | LRT $\chi^2(2)=7192.95$; $p < 0.0001$ |

**Supplementary Table 1**: Log-likelihood fit in mixture model for Experiment 1. Table 1 provides a summary of the log-likelihood fit of the 1 component, 1 parameter mixture model, the 2 component, 3 parameter mixture model, and the likelihood ratio test. Adding additional distributions had little effect as the log-likelihood was already saturated after adding 2 distributions.

| | Average Switching Probability in persistent regime | Average Switching Probability in switching regime | Average weight of both types of regimes |
|---|---|---|---|
| Mice | 0.20 ± 0.11 | 0.66 ± 0.18 | 0.67 ± 0.20 |
| Monkeys | 0.065 ± 0.038 | 0.62 ± 0.21 | 0.59 ± 0.16 |
| Humans | 0.12 ± 0.12 | 0.47 ± 0.24 | 0.67 ± 0.21 |

**Supplementary Table 2:** Average Switching Probabilities and Weight Metrics for Experiment 1. Table 2 provides a summary of the average switching probabilities in persistent and switching regime for different species, including mice, monkeys, and humans. statistics for the weight of both regimes: humans-mice: $p = 0.88$, $b = 0.003$, 95% CI = [-0.033, 0.038], $t(500) = 0.15$; humans-monkeys: $p = 0.22$, $b = -0.078$, 95% CI = [-0.20, 0.047], $t(341) = -1.22$, monkeys-mice: $p = 0.036$, $b = 0.097$, 95% CI = [0.006, 0.19], $t(341) = 2.10$

| | Average Switching Probability in persistent regime | Average Switching Probability in switching regime | Average weight of both types of regimes |
|---|---|---|---|
| Humans - 2 targets | 0.096 ± 0.082 | 0.31 ± 0.097 | 0.71 ± 0.20 |
| Humans - 3 targets | 0.12 ± 0.10 | 0.37 ± 0.094 | 0.71 ± 0.16 |
| Humans - 4 targets | 0.11 ± 0.079 | 0.40 ± 0.077 | 0.73 ± 0.16 |

**Supplementary Table 3**: Average Switching Probabilities and Weight Metrics for Experiment 2. Table 3 provides a summary of the average switching probabilities in persistent and switching regime for humans performing the task with a different number of targets.
